# Co-regulatory changes in *Homo sapiens* prefrontal cortex shape RNA-chromatin interactions

**DOI:** 10.64898/2026.03.14.711601

**Authors:** A. Vitriolo, O. Leonardi, V. Finazzi, D. Castaldi, L. Basile, F. Prazzoli, A. Lambolez, K. Yasuzawa, WH Yip, M. Murata, CW Yip, A. Karabacak, R. Pracana, R. Lehmann, JN Tegner, V. Lagani, D. Gomez-Cabrero, J. Shin, T. Kasukawa, H. Takahashi, M. Kato, M. Bienko, P. Carninci, C. Boeckx, G. Testa

## Abstract

Noncoding RNAs (ncRNAs) regulate gene expression through RNA–chromatin interactions, yet their role in human brain evolution remains unclear. To map these interactions, we integrated FANTOM6 neurogenic data with a comprehensive atlas of regulatory regions active during early cortical development. This integration effort linked ncRNAs and transcription factors (TFs) to their genomic targets, revealing functional interactions captured by RNA-DNA contact mapping, yet largely missed by chromatin conformation data. Overlaying these interactions with nearly fixed variants in *Homo sapiens* relative to extinct hominins uncovered widespread rewiring of TF binding, affecting genes implicated in progenitor proliferation and neuronal differentiation. Gene regulatory network reconstruction identified TEAD2 and ONECUT2 as key regulators of intermediate progenitors and migrating neurons. Functional perturbation followed by Cut&Tag and ATAC-seq linked the two TFs to disease- and evolutionarily- relevant pathways. Together, these results show how ncRNA–TF cooperation and selective *cis*-regulatory rewiring contributed to sapiens-specific features of cortical development.

## Main

Noncoding RNAs contribute to gene expression regulation^1–3^, yet their systematic profiling has not been possible until technologies such as RNA And DNA Interacting Complexes Ligated and sequenced (RADICL-seq) were developed^3,4^. These regulatory RNAs typically work in conjunction with transcription factors^5,6^ and this activity is often dysregulated in human disorders^7,8^. Transcription factors bind DNA by affinity, recognising nucleotide sequences that can be profiled by chromatin immunoprecipitation and analogous techniques, and modelled as binding motifs.

Despite these advances, a systematic characterization of the contribution of noncoding RNAs in modulating transcription factors activity along human brain development is missing. Moreover, we lack a reconstruction of how human-specific genetic changes reshaped transcription factor binding landscapes. We expect these changes to converge on pathways critical for neocortical development, potentially constituting risk towards or protection against human disorders. To verify this hypothesis, we integrated high-resolution maps of RNA–DNA contacts generated by RADICL-seq (Lambolez *et al.*, in preparation) with transcription factor binding profiles and single-cell multiomics, seeking for new classes of regulatory interactions that would be invisible to any of these approaches if used in isolation.

High-frequency variants (HFVs) nearly fixed in modern humans were found to substantially rewire TF binding at active cortical regulatory regions, and RADICL-seq links this rewiring to ncRNA–TF circuits controlling progenitor proliferation and neuronal differentiation, going beyond canonical TF–gene pairs and genetic regulatory changes^9–12^.

## Results

To connect nearly fixed high-frequency variants (HFVs) with transcription-factor (TF) binding changes in the developing cortex, we assembled **CatReg (see Methods)**, a curated catalogue of active cortical *cis*-regulatory elements (CREs) with TF binding support (Fig. 1A–C; Fig. S1A). CatReg integrates fetal bulk^13,14^ and single-cell chromatin accessibility with cortical organoid multi-omics^15^. Moreover, it combines motif models with orthogonal genome-binding evidence from ChIP-seq/CUT&Tag resources^16^ (see Methods). In total, CatReg comprises ∼250,000 CREs linked to putative target genes (Fig. 1B) and includes chromatin-associated regulators beyond sequence-specific DNA-binding factors (Fig. 1C).

**Figure 1.**
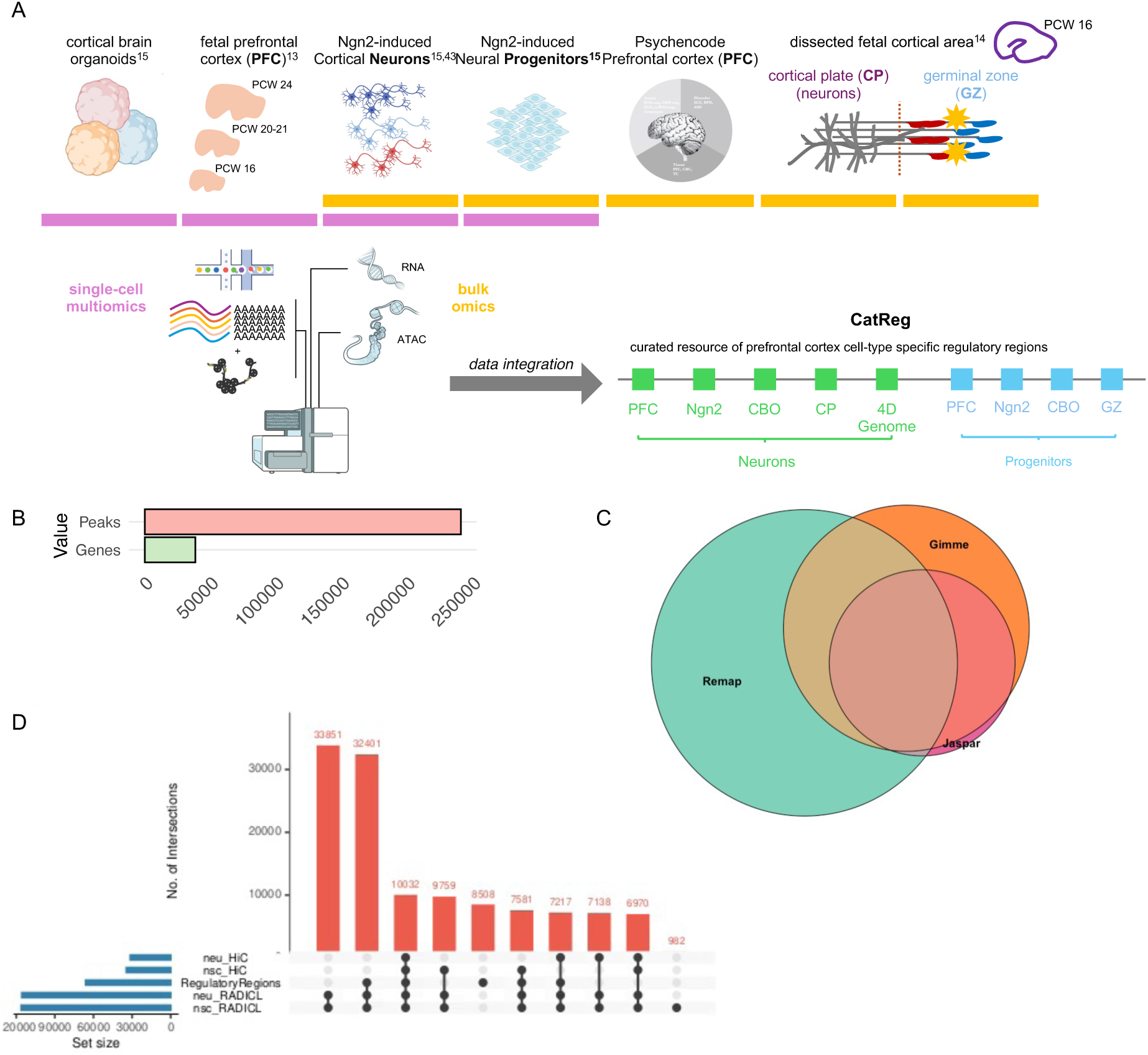
Building a regulatory landscape of early cortical development. **A.** Schematic of the multi-omics datasets integrated to construct CatReg, a catalogue of cortical cis-regulatory elements (CREs) active during early development across progenitor and neuronal states (illustrated developmental stages in post-conception weeks, PCW). **B.** Size of the CatReg resource, showing the number of CREs and associated genes. **C.** Overlap between TF resources used to assign regulator support to CatReg CREs, integrating motif collections (JASPAR and GimmeMotifs) with orthogonal TF binding evidence aggregated from ReMap; circle area is proportional to the number of regulators represented in each resource. **C.** UpSet plot showing intersections between CatReg CREs and RADICL-seq or Hi-C contacts in NSCs and neurons.

A key advantage of incorporating experimental binding data is that it captures regulators that engage chromatin indirectly—for example, proteins that interact with histones, participate in transcriptional complexes, or modulate DNA-binding factors. We therefore retained these chromatin-associated regulators in the set of TFs considered here, given their physical association with the genome and their potential contribution to transcriptional control.

To confirm that CatReg captures active regulatory regions, we integrated independent FANTOM6 datasets (RADICL-seq, Hi-C and CUT&Tag; Fig. S1B–D) and verified that CatReg regions are enriched for active enhancer (H3K4me1, H3K27ac) and promoter (H3K4me3) marks in progenitors and neurons. We then compared the overlap of CatReg with chromatin contacts (Hi-C) and RNA–DNA contacts (RADICL-seq) in both progenitors (NSC) and neurons (Neu). Hi-C interactions spanned broader genomic distances (median ∼500 kb; Fig. S1E), whereas RADICL-seq interactions were more proximal (median <200 kb; Fig. S1F). Consistent with this, Hi-C contacts showed a lower overlap with CatReg, suggesting that in human neural lineages Hi-C does not preferentially capture functional regulatory interactions, as observed in mouse embryonic stem cells^17^. By contrast, RADICL-seq contacts overlapped CatReg more extensively (Fig. 1D), supporting the regulatory relevance of RNA–DNA interactions.

To refine *cis*-regulatory links, we inferred topologically associated domain (TAD) boundaries using a motif-orientation strategy based on convergent CTCF motifs^18^ by adapting the original approach from Nanni et al.^18^, which leverages accessible regions and co-accessibility maps that can be drawn from single-cell ATAC data by repurposing these maps as a surrogate of contact matrices (see Methods, Fig. S1G). Restricting interactions to those occurring within inferred TADs yielded *cis*-regulatory distance distributions consistent with micro-C data from human and mouse embryonic stem cells^19,20^ (Fig.S1H). Finally, intersecting CatReg with ReMap ChIP-seq data revealed a non-linear relationship between the number of regions bound by a given TF and the number of associated target genes (Fig. S1I).

Together, these analyses indicate that CatReg enriches functional TF–gene regulatory associations, highlighting **993 regulators** with supported binding evidence (Fig. S1J; hereafter referred to as TFs).

To dissect how RNA–DNA interactions may have contributed to early neocortical development in *Homo sapiens*, we focused on genomic sites carrying evolutionarily relevant variation. We leveraged a genome-wide set of high-frequency variants (HFVs)^21^, identified by comparing modern human genomes with Neanderthal and Denisovan sequences, retaining modern human–specific alleles present in an estimated >90% of present-day individuals. We then appraised the impact of HFVs on transcription factor (TF) binding affinity using motifbreakR, restricting analyses to variants classified as “strong” disruptions (Fig. 2A; Fig. S2A). TFs were prioritized by the number of affected binding sites (Fig. 2B) and independently by the magnitude of predicted affinity change (Fig. 2C; Fig. S2B).

**Figure 2.**
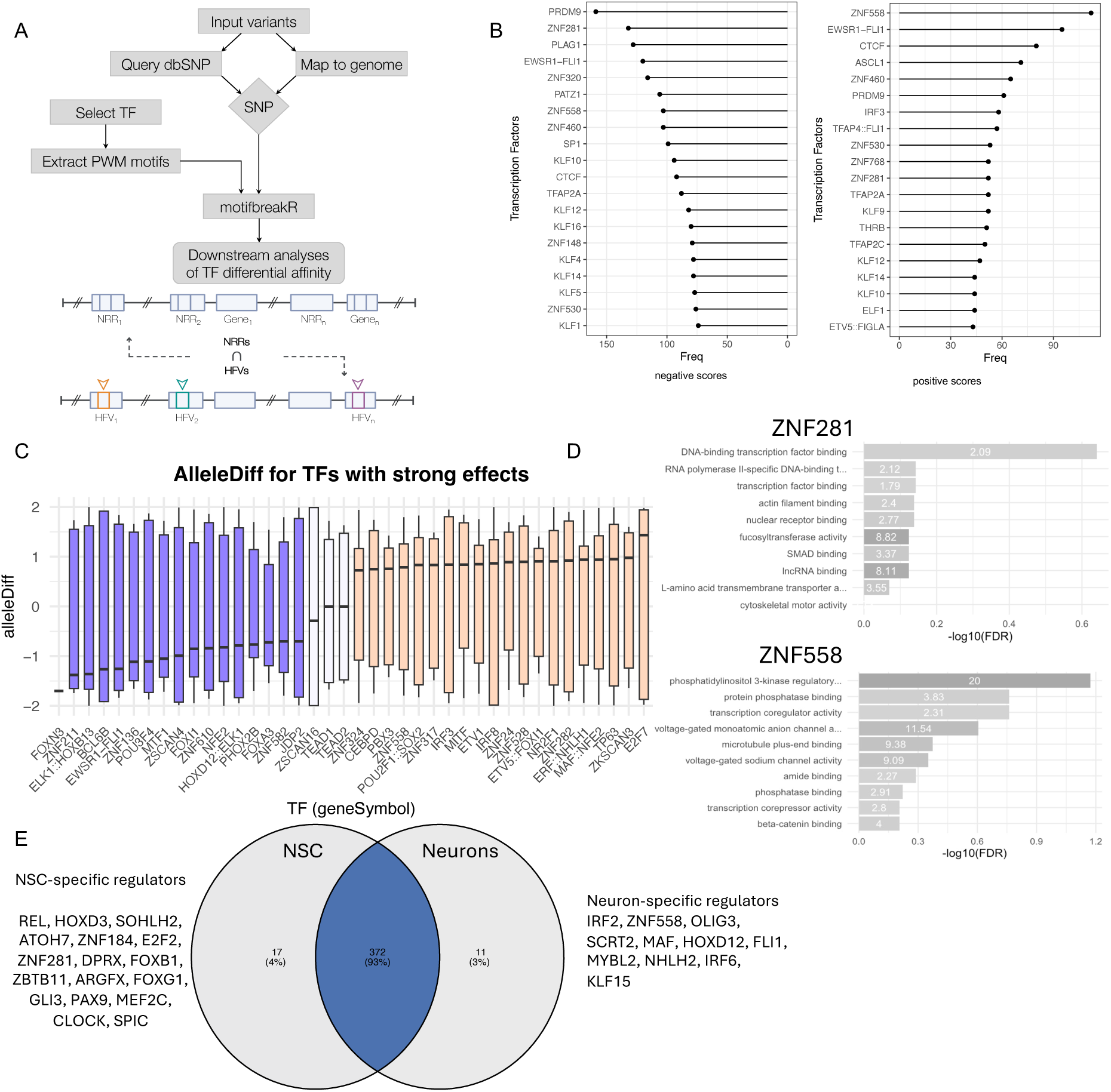
Modern human–specific high-frequency variants reshape predicted TF binding. **A.** Workflow to quantify nucleotide-resolution changes in TF motif affinity caused by high-frequency variants (HFVs) using motifbreakR and JASPAR 2024 PWMs, and to propagate predicted effects to downstream regulatory analyses. **B.** TFs ranked by the number of CatReg sites containing HFVs predicted to cause **strong** decreases (left) or increases (right) in motif affinity (AlleleDiff). **C.** Distribution of predicted affinity changes (AlleleDiff) for top affected TFs. TFs are grouped by median AlleleDiff: predominantly negative (left), near-zero (|median| ≤ 0.5; middle), or predominantly positive (right). **D.** GO Biological Process enrichment for inferred targets linked to ZNF281- or ZNF558-associated CREs whose motifs are disrupted by HFVs. Bars show –log10(FDR). **E.** Overlap of HFV-affected regulators in neural stem cells (NSCs) and neurons whose target genes are also contacted by ncRNAs in RADICL-seq (FANTOM6); NSC- and neuron-enriched regulators are listed.

Across both rankings, the chromatin organizer CTCF emerged among the most affected factors for both increased and decreased predicted affinity, consistent with widespread human-specific redistribution of CTCF binding motifs and potential changes in higher-order chromatin architecture between modern and archaic humans. Zinc Finger Protein 281 (ZNF281), Zinc Finger Protein 460 (ZNF460), Krüppel-like factor 10 (KLF10), Transcription Factor AP-2 Alpha (TFAP2A) and PR/SET domain containing protein 9 (PRDM9) showed similarly pervasive bidirectional shifts. PRDM9, best known for its role in meiotic recombination hotspot specification via H3K4 trimethylation^22^, also appeared among the top-ranked motif models. This either raises the possibility that HFVs also contribute to broader changes in promoter and regulatory activity landscapes or potentially highlights a function for PRDM9 in the cortex; both hypotheses will require future validation. The top-ranked motif models additionally include the JASPAR entry annotated as EWSR1–FLI1, which represents a composite/fusion-associated binding model. Because predicted disruptions to this motif could reflect sequence features attributable to either one component TF, both, or to the composite model itself, this signal must be taken as hypothesis-generating rather than evidence of fusion-specific regulation in cortex.

Effect-size ranking revealed predominantly bidirectional shifts in predicted affinity across top factors, suggesting extensive rewiring of TF binding by the large number of modern human–specific variants considered. Only three TFs—ZSCAN16, TEAD1 and TEAD2—showed a median differential affinity close to zero, consistent with a near balance of predicted gains and losses, whereas FOXN3 showed a smaller but consistently negative shift.

Two zinc-finger TFs, ZNF281 and ZNF558, were prioritized by both ranking strategies. Given their limited functional characterization in this context, we performed GO enrichment on their inferred target genes to contextualize pathways most affected by HFVs (Fig. 2D). Targets linked to ZNF281-associated HFV-disrupted CREs were enriched for RNA-related functions, alongside terms related to actin cytoskeleton and SMAD-associated signalling. These enrichments are consistent with prior reports implicating ZNF281 in RNA-mediated regulation^23^ and SMAD modulation^24^ and with roles for cytoskeletal regulation during cell-cycle dynamics in neural progenitors^25^. ZNF281 has also been implicated in regulatory variation in neural precursors, including effects attributed to modern human–specific variants^26^ In contrast, ZNF558-linked targets were enriched for transcriptional regulation and post-translational modification, and included terms related to microtubule organization and voltage-gated sodium channel activity; ZNF558 expression has been reported in human, but not chimpanzee, forebrain neural progenitors^27^ and has been associated with psychiatric phenotypes^28^.

Finally, to connect HFV-affected TF programs to RNA–DNA regulation, we integrated RADICL-seq data from neural stem cells (NSCs) and neurons (Lambolez et al. this collection) to identify TFs whose target genes are also contacted by non-coding RNAs (Fig. 2E). Most ncRNAs participating in these TF–RNA–DNA triplets were shared across NSCs and induced glutamatergic neurons, although a subset was cell type–specific. Notably, FOXG1 and GLI3, which are key regulators of radial glia identity and cortical evolution^12^, were among the TFs participating in these evolutionarily relevant interaction triplets.

Building on CatReg and the RADICL-seq–derived ncRNA–gene contact map, we next asked how ncRNA–TF cooperation contributes to establishing cell identity during human corticogenesis, and whether this layer preferentially involves TFs whose binding landscapes were reshaped by **modern human–specific regulatory changes captured by high-frequency variants (HFVs)**. To this end, we integrated single-cell transcriptomic trajectories with our TF- and ncRNA-centered interaction resources to identify cell type–specific TF–ncRNA–gene triplets that operate during neurogenesis.

We reconstructed a gene regulatory framework for fetal prefrontal cortex (PFC) by re-analysing the cortical arealization atlas^29^ and focusing on PFC cells (Fig. 3A). After integration and transfer of refined cell annotations^30^ (see Methods), this dataset distinguishes radial glia subtypes, intermediate progenitors (IP), and successive excitatory neuronal states, including early migrating excitatory neurons (Exc_Mig). To infer cell type–specific GRNs while explicitly incorporating ncRNAs as regulatory inputs, we used CellOracle^31^, which supports integration of external interaction priors and enables forward simulation of network responses to in silico perturbations (see Methods). We introduced RADICL-seq–derived ncRNA–gene contacts as an additional layer of directed regulatory evidence, allowing ncRNAs to act as candidate upstream regulators alongside TFs, rather than appearing only as downstream targets. We then prioritized regulators that best explain cell identity by intersecting regulators with strong network influence with cell type marker programs, yielding ∼200 candidate master regulators (Fig. 3B; Fig. S4B). Ranking these regulators by their contribution to controlling the top marker genes in each cell type highlighted a small set of highly stage-restricted factors, including TEAD2 in IP, ONECUT2 in inhibitory neurons (InhN), DBX2 in outer radial glia (oRG), and GLI3 in ventral radial glia (vRG) (Fig. 3C). Relaxing thresholds expanded this set while preserving cell type–specific expression patterns (Fig. S4C–D).

**Figure 3.**
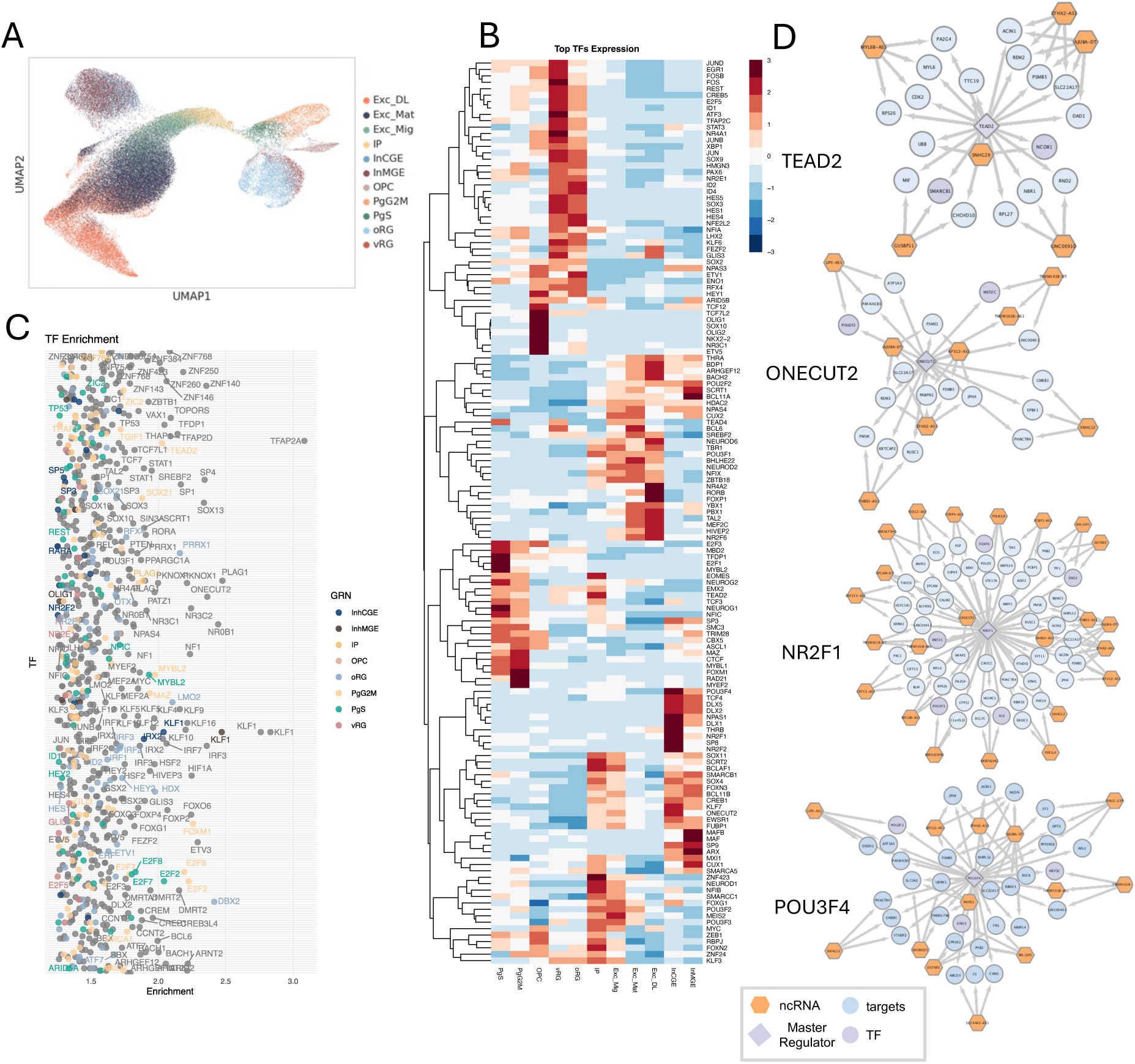
Modern human regulatory signals converge on TF–ncRNA networks during neuronal fate acquisition. **A.** UMAP of fetal prefrontal cortex (PFC) single-cell RNA-seq data reanalysed from Bhaduri *et al.*^29^ Cell types include ventral radial glia (vRG), outer radial glia (oRG), cycling progenitors (PgS, PgG2M), intermediate progenitors (IP), oligodendrocyte precursors (OPC), inhibitory interneuron lineages (InhN) from the medial and caudal ganglionic eminences (InMGE, InCGE respectively), and excitatory neuronal states (Exc_Mig, Exc_Mat, Exc_DL). **B.** Expression of prioritized transcription factors across PFC cell types (pseudobulk log-normalized counts; row-wise z-scores), highlighting cell type–restricted regulators. **C.** Cell type–specific regulator enrichment scores, ranking TFs by their contribution to cell identity programs. For each TF, enrichment reflects the fraction of cell type marker genes among its inferred targets (colored by the cell type in which the TF is prioritized). **D.** TF-centered subnetworks integrating cell type–specific GRNs with RADICL-seq ncRNA–gene contacts (examples: TEAD2, ONECUT2, NR2F1, POU3F4). Diamonds denote TF hubs; hexagons denote ncRNAs; circles denote target genes.

In parallel, we filtered the RADICL-seq interactome in neural stem cells (NSCs) to prioritize reproducible ncRNA–gene contacts and reduce sparsity-driven links (Fig. S5A). To focus on ncRNAs with master regulatory activity, we kept only interactomes of ncRNAs sharing at least two targets with other ncRNAs (Fig. S5B), we quantified the number of targets they shared with TFs/master regulators previously identified (Fig. S5C). Genes jointly targeted by ncRNAs and cell type marker regulators showed a significant overlap with targets of TFs predicted to display **HFV-associated differential binding between the modern human and ancestral allelic states** (Fig. S5D). Extending the analysis from marker regulators to the full GRN did not increase this overlap (Fig. S5E), indicating that—at least in progenitors—ncRNA–TF co-regulation preferentially involves cell identity–defining TFs rather than broadly expressed regulators. Based on this enrichment, we expanded triplet discovery by allowing ncRNAs that share at least one target with HFV-affected TFs.

The combination of developmentally informed cell type-specific TF prioritization, with enriched nc-RNA co-regulatory activity, revealed a TF-ncRNA convergent activity centered on the IP→early neuron transition. Indeed, although ONECUT2 was prioritized as an InhN regulator, it is also prominently expressed in Exc_Mig, whereas TEAD2 is most active in IP, which precede Exc_Mig along the reconstructed trajectory. We therefore focused on TF–ncRNA–gene triplets associated with these transitions and extracted neuroblast/NPC subnetworks for TEAD2 and ONECUT2 (Fig. 3D; Fig. S5F). We also extracted subnetworks for increased binder POU3F4, which binds hundreds of regions affecting differential expression between *Homo* groups^32^, and decreased binder NR2F1, which show among the strongest HFV-associated differential affinity shifts within the master-regulator (Fig. 3D; Fig. S5F). Notably, pseudotime analysis indicates that these four TFs peak immediately before neuronal differentiation (Fig. S4E), and cells at these peak coordinates are dominated by IP and Exc_Mig states. Together, these results place **modern human HFV-associated TF binding differences** and ncRNA–TF co-regulation at the developmental interval where progenitors commit to neuronal identity.

To functionally elucidate the regulatory programs coordinated by transcription factors and ncRNAs during cortical neurogenesis, we examined the gene sets jointly targeted by TF–ncRNA pairs across progenitor and neuronal populations. Shared targets in intermediate progenitors (IP) were enriched for processes controlling cell-cycle progression, mitotic transition and protein turnover, whereas shared neuronal targets were associated with chromatin organization, cytoskeletal assembly and synaptic maturation (Fig. 4A). In line with these stage-specific programs, TF interactomes overlapped more extensively in progenitors than in neurons (Fig. 4B), indicating that regulatory cooperation is broad early and becomes progressively refined as cells commit to mature lineages.

**Figure 4.**
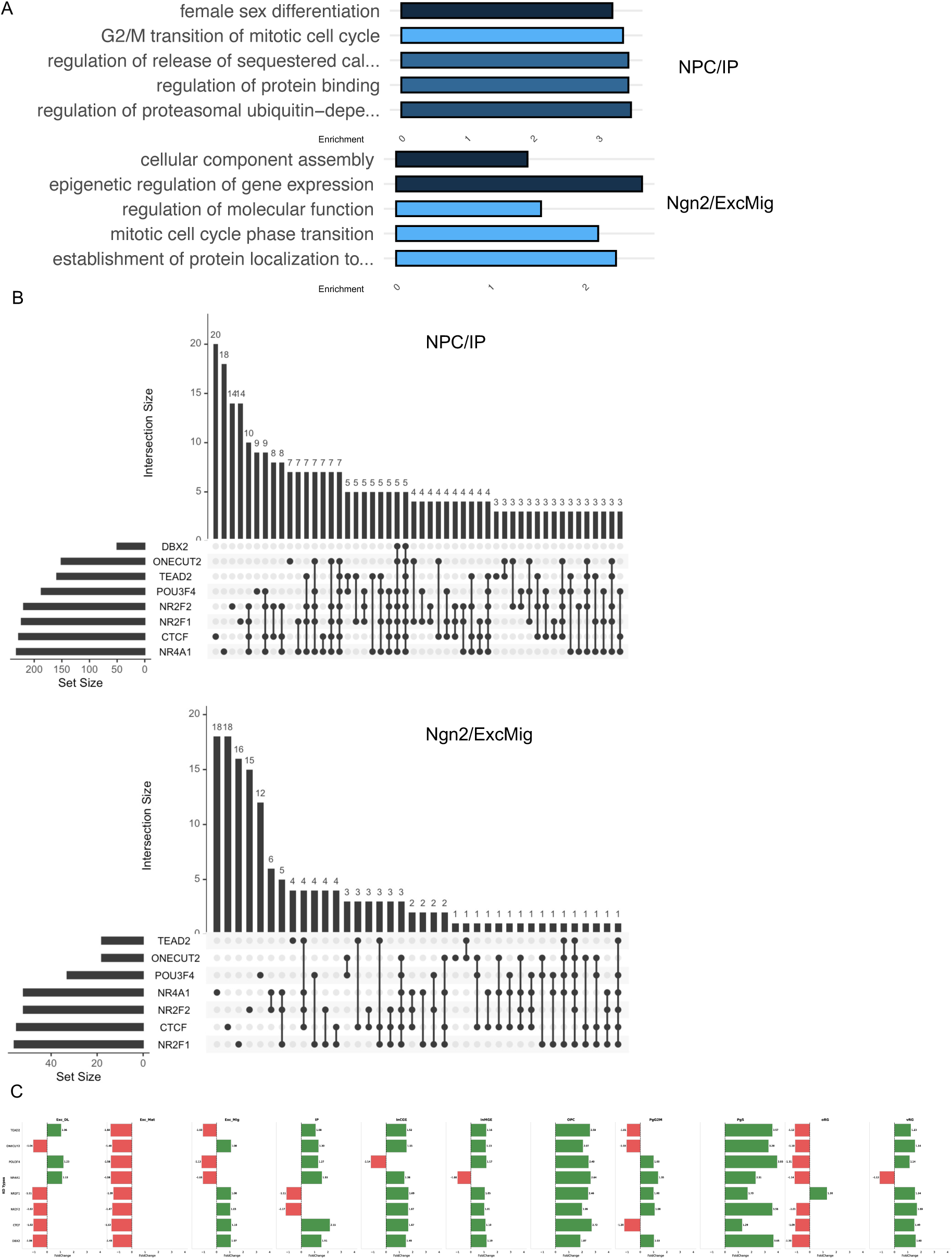
Functional consequences of HFV-affected TF–ncRNA regulatory programs during cortical neurogenesis. **A.** Gene Ontology (GO) enrichment of genes jointly targeted by consistent TF–ncRNA pairs in progenitor (NPC/IP) and early neuronal (Ngn2/ExcMig) populations. **B.** UpSet plots summarizing overlap among target-gene sets of prioritized core TFs within progenitor (NPC/IP) and early neuronal (Ngn2/ExcMig) populations. **C.** CellOracle virtual knock-out (KO) simulations for core TFs. For each KO (one facet per TF), bars report the log2 fold-change (log2FC) in the proportion of each annotated cell type in the KO relative to the unperturbed reference (log2FC > 0 indicates expansion; log2FC < 0 indicates depletion). Negative values are reported in red, while positive log2FC are reported in green.

We next leveraged CellOracle to infer the developmental consequences of perturbing TFs prioritized from the HFV-focused analysis. Virtual knock-out of TEAD2, ONECUT2, POU3F4 or NR2F1 disrupted neurogenic progression and reduced the abundance of outer radial glia (oRG) and mature excitatory neuronal states (Fig. 4C). Across perturbations, the strongest and most consistent effect was a depletion of oRG, linking these HFV-affected regulators to a progenitor population implicated in primate-specific cortical expansion^33–35^. oRGs arise at later developmental stages and expand markedly in primates^36,37^, a process thought to depend on the tuning of cell-cycle dynamics and mitotic spindle orientation.

Comparing perturbation phenotypes further resolved two response modes. ONECUT2 knock-out decreased both deep-layer (Exc_DL) and mature (Exc_Mat) excitatory neurons, whereas TEAD2 knock-out increased Exc_DL while decreasing Exc_Mat (Fig. 4C). Earlier along the trajectory, ONECUT2 knock-out increased Exc_Mig and IP, whereas TEAD2 knock-out decreased Exc_Mig while increasing IP. Despite these differences in excitatory-state balance, TEAD2 and ONECUT2 converged in their effects on progenitor composition, producing similar shifts when comparing oRG relative to vRG, and both perturbations increased oligodendrocyte precursors (OPCs). Together, these results implicate TEAD2 and ONECUT2 as regulators of the progenitor-to-neuron transition that shape the acquisition of IP and oRG identities—two cell states central to the expansion of the human neocortex.

To experimentally validate the regulatory divergence suggested by our virtual perturbations, we performed targeted knock-down (KD) experiments for two HFV-affected regulators with contrasting predicted effects on neurogenic trajectories: TEAD2 and ONECUT2. Specifically, we carried out TEAD2 KD in human iPSCs and ONECUT2 KD in cortical neurons, and profiled chromatin accessibility changes at cis-regulatory elements (CREs) (see Methods). Integrating these datasets with FANTOM resources enabled systematic mapping of TF-bound CREs to downstream gene programs and phenotypic annotations. KD of the two TFs elicited distinct chromatin accessibility responses at their bound CREs. TEAD2-bound CREs that became differentially accessible upon KD were linked to genes involved in cell migration and protein transport, whereas ONECUT2-bound CREs that changed upon KD were enriched for synaptic function and neuronal morphogenesis (Fig. 5A), consistent with a progenitor-oriented versus neuron-maturation regulatory bias.

**Figure 5.**
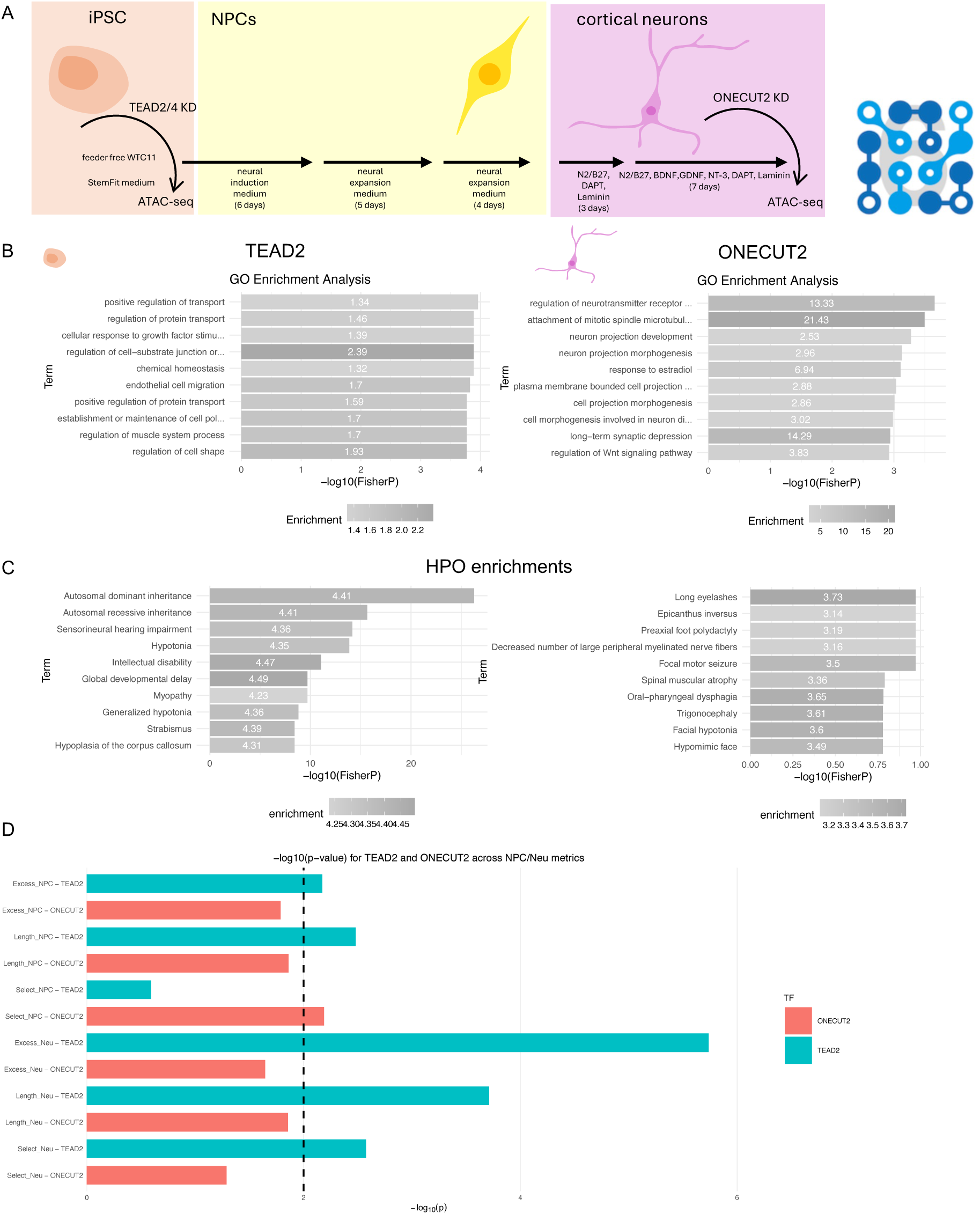
Experimental validation distinguishes a progenitor-linked TEAD2 module from a neuronal ONECUT2 module. **A.** Functional enrichment of genes associated with TF-bound cis-regulatory elements (CREs) that change chromatin accessibility upon knock-down (KD). TEAD2 KD was performed in human iPSCs and ONECUT2 KD in cortical neurons; enrichments were computed from genes linked to differentially accessible, TF-targeted CREs. B. Gene Ontology (GO) enrichment of genes associated with KD-responsive TF-targeted CREs, highlighting distinct phenotype classes linked to ONECUT2- versus TEAD2-regulated programs. **C.** Human Phenotype Ontology (HPO) enrichment of genes associated with KD-responsive TF-targeted CREs, highlighting distinct phenotype classes linked to ONECUT2- versus TEAD2-regulated programs. **D.** Enrichment of evolutionary annotations among genes linked to KD-responsive TF-targeted CREs. TEAD2-associated targets show increased overlap with regions carrying human-lineage regulatory signatures (e.g., human-specific substitutions, elevated mutation-rate metrics, and positive selection signals as defined in Methods), whereas ONECUT2-associated targets show comparatively limited overlap.

Human Phenotype Ontology (HPO) enrichment of genes associated with KD-responsive regulatory regions further supported this functional split. ONECUT2-linked CREs were enriched for phenotypes related to cognitive and synaptic dysfunction, whereas TEAD2-linked CREs were enriched for neurodevelopmental and structural brain phenotypes (Fig. 5B), consistent with modulation of progenitor behaviours and cortical growth. Given evidence that expansion of radial glia lineages contributes to increased cortical surface and gyrification in humans^38,39^, these results support a model in which ONECUT2 primarily tunes neuronal differentiation and maturation programs, whereas TEAD2 preferentially impacts progenitor-linked processes relevant to cortical architecture.

Finally, integrating these functional datasets with evolutionary annotations revealed that TEAD2-regulated genes are preferentially located in genomic regions enriched for human-specific substitutions, excessive mutation rates, and signatures of positive selection, whereas ONECUT2 targets showed comparatively limited overlap with such features (Fig. 5C), suggesting a more specific and confined impact during evolution. Together, these results delineate two complementary axes of cortical regulation: a neuronal differentiation module associated with ONECUT2, and a progenitor-linked module associated with TEAD2 that is disproportionately connected to genomic regions bearing signatures of recent human evolution.

## Conclusions

In this work, we integrated RNA–chromatin interaction maps with multi-layer regulatory genomics to extend the FANTOM6 framework and define how noncoding RNAs and transcription factors cooperate during human cortical development. By combining RADICL-seq with bulk and single-cell omics, we generated a cortex-focused catalogue of active regulatory elements and RNA-linked connectivity across neurogenesis. This integrative strategy connects physical RNA–DNA contacts to functional enhancer–promoter regulation, capturing a regulatory dimension that is not resolved by chromatin conformation assays alone.

Placing these regulatory maps in an evolutionary context, we show that modern HFVs preferentially reshape transcription factor binding landscapes at regulators implicated in neurogenesis and cortical expansion. The involvement of POU3F4 and NR2F1 at human-derived regulatory sites points to species-specific remodeling of transcriptional programs and to regulatory mechanisms relevant to rostro–caudal patterning. In parallel, the widespread signatures across CTCF, PRDM9 and multiple KRAB-ZFP family members suggest coordinated tuning of chromatin organization and local transcriptional circuits in the human lineage. Within this set, ZNF281 and ZNF558 emerge as candidate mediators of RNA-linked chromatin regulation, connecting transcriptional control to actin cytoskeletal remodeling—processes repeatedly implicated in human brain evolution^40–43^.

Integrating RADICL-seq contacts into cell type–specific gene regulatory networks revealed a core set of TF–ncRNA regulatory modules that track progenitor diversification and neuronal commitment. Evolutionarily sensitive regulators such as TEAD2 and ONECUT2 occupy central positions at the transition from intermediate progenitors to migrating and maturing neurons. Virtual perturbations predicted that both factors are required for efficient production of outer radial glia, a progenitor population associated with primate cortical expansion. Targeted knockdown followed by ATAC-seq further supports functional specialization of these regulators: ONECUT2-linked regulatory elements preferentially connect to neuronal morphogenesis and synaptic programs, whereas TEAD2-linked elements connect to migration and protein transport pathways. Notably, TEAD2-associated targets show stronger overlap with genomic regions carrying human-lineage evolutionary signatures, consistent with a disproportionate involvement in recently modified regulatory circuitry.

Together, our results support an integrated regulatory logic in which noncoding RNAs help organize transcription factor activity at selected genomic sites, shaping gene programs that underlie key transitions in cortical development. More broadly, this work provides a blueprint for coupling RNA–DNA connectivity with multi-omics and evolutionary genomics to infer the functional grammar of cis-regulation. Extending this framework across additional cell states, developmental windows and primate lineages should help resolve how regulatory innovation—and RNA-mediated modulation—has contributed to the emergence of human-specific aspects of brain development.

## Acknowledgements

Sequencing was performed by Laboratory for Genotyping Development in RIKEN IMS and the National facilities (formerly the sequencing facility) and the Centre of Genomics in Human Technopole (HT). Hi-C was sequenced at Karolinska Institutet, Stockholm.

We thank Nobuyuki Takeda and Teruaki Kitakura (RIKEN) for their support of the IT infrastructure for the FANTOM6 collaboration, and Emi Ito (RIKEN) for her administrative support.

This is a collaborative work with FANTOM6 Consortium. We thank all consortium members for their insights and suggestions. We thank present and former members of G.T.’s laboratory for insightful feedback provided on this study.

We thank the European School of Molecular Medicine (SEMM) at which O.L was enrolled as student for his PhD degree program in Systems Medicine. Some components of the schematics in Figure 1 were adapted from BioRender. Figure 1 contains NIAID Visual & Medical Arts. (10/7/2024). Next Gen Sequencer. NIAID NIH BIOART Source. bioart.niaid.nih.gov/bioart/386; bioart.niaid.nih.gov/bioart/125; bioart.niaid.nih.gov/bioart/77. C. B. was supported by Project PID2023-146627NB-I00 (Spanish Ministry of Science, Innovation, and Universities CIENCIA/AEI), project 2021-SGR-313 (AGAUR/Generalitat de Catalunya) and a Leonardo Grant for Researchers and Cultural Creators, BBVA Foundation. A.V. was supported by Telethon Research Grant GGP19226 (GT) and RE-MEND (101057604).

FANTOM was supported by research grant for the RIKEN Center for Integrative Medical Sciences (IMS) from the Ministry of Education, Culture, Sports, Science and Technology (MEXT) Japan.

This work was supported by the Human Technopole Foundation, a research foundation funded by the Italian Government under the Ministries of Economy & Finance, Health, and Education, University and Research.

## Authors contributions

A.V., C.B. and G.T. conceived this work and wrote the paper; A.V., O.L. and V.F performed most of coding and analytical activities presented; D.C. curated the single-cell RNA-seq data; L.B and F.P. contributed to the downstream single-cell analytical reconstruction of gene regulatory networks and *in silico* perturbations; A.L. performed peak calling and data analysis of CUT&Tag and Hi-C data; CW.Y., M.M., A.L, A.K.C., R.P. supported the study and provided insightful suggestions; K.Y contributed with KD-ATAC experiments in J.S. lab; WH.Y performed Hi-C experiments in M.B. lab; CW.Y., T.K., H.T., M.K. and P.C supervised the FANTOM6 project; R.L. and V.L. from JN.T. and D.G-C. labs contributed with methodological discussions and provided feedbacks on the manuscript draft. C.B., M.B., P.C., J.S., G.T. and A.V supervised the project.

## Material and Methods

All human samples used in this study were commercially available, obtained from public collections, or collected from patients under informed consent and in accordance with the recommendations from the Declaration of Helsinki. The use of all non-exempt human materials for research in this project has been approved by the RIKEN Yokohama Branch Ethics Committee (approval number RIKEN-Y-2024-098).

Induced pluripotent stem cells referred to as “iPSC” in the FANTOM6 consortium data (Hi-C, RADICL-seq and Cut&Tag used in this study) correspond to human i3N iPSCs in the male WTC11 background, harbouring a doxycycline-inducible Neurogenin 2 (NEUROG2/<u>Ngn2</u>) transgene integrated at the AAVS1 locus (C. Wang et al., 2017). These cells were kindly gifted by Dr. Michael Ward (National Institutes of Health, United States) to the consortium. Neural stem cells (NSC) and glutamatergic cortical neurons (Ngn2) were derived from this line. Single-cell multiomic data for Cortical Brain Organoids and Ngn2 cortical neurons come from the cohort of Testa group iPSC control lines described in Pereira, Finazzi et al^15^.

RNA And DNA Interacting Complexes Ligated and sequenced (RADICL-seq) was performed on all samples exactly as described in Bonetti et al., and Shu et al.^44,45^ with two to three biological replicates per sample.

### CUT&Tag library preparation

iPS, NSC and Neuron cells were harvested in two biological replicates per sample. Nuclei isolation was performed as instructed in the “Nuclei Isolation for Single CellATAC Sequencing” protocol from 10x Genomics® (Demonstrated Protocol CG000169 Rev D) after optimizing the concentration of BSA in PBS to 0.1% instead of 0.04%. The isolated nuclei were used for library preparation with the CUT&Tag-IT™ Assay Kit (Active motif®, catalog no. 53160), following the manufacturer’s instructions (Document 2200 version B6). In brief, nuclei were immobilized on concanavalin A-coated beads and magnetically separated. The nuclei were then resuspended in a buffer containing a protease inhibitor cocktail and digitonin, and 1 µg of either of the following antibodies was added to each sample (See Lambolez et al. in this collection for the complete list of antibodies, and manufacturers’ identifiers). Nuclei were left to incubate with the antibodies overnight, then once again magnetically separated, incubated for 1 hour with 1:100 guinea pig anti-rabbit secondary antibody in Dig-Wash buffer at room temperature, and finally washed several times with Dig-Wash buffer. The antibody-bound chromatin was next incubated for 1 hour with 1:100 diluted CUT&Tag-IT™ Assembled pA-Tn5 transposomes in Dig-300 buffer at room temperature, then stringently washed several times with Dig-300 buffer. The chromatin was sheared and sequence adaptors were added by tagmentation for 1 hour at 37°C. The samples were de-crosslinked for 1 hour at 55°C with buffers containing 0.1% SDS, 16 mM EDTA, and 1 µg proteinase K, which was followed by DNA extraction. Libraries were amplified by PCR with the index primers, and size selection was performed using solid phase reversible immobilization (SPRI) beads to remove primer dimers. Sequencing was performed with a HiSeq X™ Ten sequencing system (Illumina®) in the following conditions: R1, 150 cycles; R2, 150 cycles; Index1, 8 cycles; Index2, 8 cycles.

### CUT&Tag data processing

Raw sequencing data from CUT&Tag were trimmed, aligned to the hg38 genome, sorted and deduplicated using the ENCODE ATAC-seq pipeline version 1.7.0 (D. S. Kim, 2023), with default parameters.

### Hi-C library preparation

iPSC, NSC and Neuron samples were cross-linked by exposure to 1× PBS (Wako®, FUJIFILM®, catalog no. 166-23555) solution supplemented with 2% formaldehyde (Thermo Fisher Scientific®, catalog no. 28906) for 10 min at room temperature under rotation. The formaldehyde was then quenched and washed, and cells were pelleted at 200 *g* for the iPSC sample, 300 *g* for the NSC sample and 150-200 *g* for the Neuron sample, before storage at-80°C. In situ proximity ligation and Hi-C library preparation were performed with the Arima-HiC+ kit (Arima Genomics®, catalog no. PNA510008) and the Arima Library Prep Module (ArimaGenomics®, catalog no. PNA303011) following the manufacturer’s instructions (Document A160134 v01). Library were sequenced on an Illumina® NovaSeq™6000 platform using the two lanes of the S4 Reagent Kit v1.5 (300 cycles, Illumina®, catalog no. 20028312) in paired-end 150 sequencing mode.

### Regulatory Regions collection generation and genomic data curation

The catalog of regulatory regions (CatReg) was compiled through the integration of different epigenomic data at peak level. Prefrontal cortex data from Psychencode was gathered from the psychencode.org website (DER-04a_hg38lft_PEC_enhancers). This data was combined with open chromatin regions from post-conception week 15–17 human neocortex^14^. We expanded this list with single-cell data from Ngn2-induced 2D neural series, and cortical brain organoids data from ^15,46,47^, using co-accessible chromatin regions derived from single-cell multiomics data (see E-MTAB-16422 and associated entries, GEX/ATAC multiomics). Contacts or peaks for Hi-C, RADICL-seq, ChIP-seq and CUT&Tag of histone post-translational modifications, as well as CUT&Tag data for ONECUT2 and TEAD2/4, and ATAC-seq differentially available sites upon KD of the two TF were generated by the FANTOM6 consortium (Lambolez *et al.*, in preparation). Peak-gene association was derived from each individual study. Genome binding protein data was derived from the Remap2022^16^ website (https://remap2022.univ-amu.fr/) by downloading the full list annotated on hg38 genome coordinates. CTCF motif location was derived from PWMScan ^48^. Significance of overlap between CatReg and RADICL-seq interaction maps, or Hi-C contact maps was performed via hypergeometric test. When bedtools fisher or R hypergeometric tests (phyper) generated close to zero significance values, significance of peak set overlaps was assessed using an empirical permutation test. Briefly, peaks of one set (CatReg) were randomly relocated (N=5,000 permutations), preserving interval lengths and chromosome assignment (constrained to a genome mask removing ENCODE exclusion regions). Overlap with the test set (e.g. RADICL- or Hi-C contacts) was quantified as intersecting base pairs, and empirical p-values were computed as the fraction of permutations with overlap equal to or greater than observed.

### TAD boundaries inference in single-cell ATAC data

Following Nanni et al’s work on CTCF spatial patterning^18^, we developed a simple method to leverage CTCF motif orientation at putative CREs found by means of co-accessibility, which was calculated from single-cell data in a cell-type specific fashion^15^. Single-cell data was gathered from control lines in E-MTAB-16422. We used progenitors and neurons inferred TAD boundaries to refine the list of regulatory regions in our CatReg. Code is available at https://github.com/veronica-fin/TADboundaries

### Quantification of variants impact on transcription factors binding sites

To quantify the impact of sapiens high frequency variants on genome binding we adapted the protocol from Moriano et al. ^49^ to work with the newly available JASPAR 2024 dataset^50^. Briefly, to evaluate disruptions of TFBS, we generated a set of genomic coordinates of variants using a unique identifier based on genomic coordinates and allele information. Differences in TF binding affinity were computed, applying motifbreakR^51^ and using position weighted matrices annotated in the JASPAR2024 motif collection. A significance threshold was set to 1e-5, and an even background nucleotide distribution was assumed. We performed multiple test correction through Benjamini-Hochberg method. Redundant motifs were dropped. TF-variant associations were further filtered based on predicted affinity difference above the fourth quantile of the distribution.

### Adaptation of single-cell RNA-seq from Bhaduri et al ^29^

In order to integrate CatReg with transcriptomics data, we chose to adapt prefrontal cortex single-cell RNA-seq data from Bhaduri et al. First, we extracted PFC-specific cells, and consequently ingested this data into previously published single-cell data^30^ which provided a more detailed cell-type annotation, distinguishing progenitors and neurons classes. We used Scanpy 1.9.2 to normalize counts, perform PCA, and generate a kNN graph, using the first 8 principal components, and k=100 cells per neighborhood. Based on this kNN graph we generated a UMAP, and leiden clusters, using resolution=0.8, to verify that cell type annotations ingested from Polioudakis et al^30^ were consistent. After, we calculated diffusion pseudotime^52^, diffusion maps^53^, we applied PAGA^54^ and Drawgraph to cluster cells along putative differentiation trajectories, using ForceAtlas2 algorithm^55^.

### Gene regulatory network reconstruction

To build cell type specific gene regulatory networks (GRN) we leveraged CellOracle^31^, combining CatReg as an annotated list of regulatory regions associated to genes, and Bhaduri et al. PFC single-cell RNA-seq data, as a transcriptomic layer. Additional TF-gene associations derived from Remap were added after generation of the base GRN. CellOracle was also used for virtual KO experiments. Differential cell type abundances upon KO were measured by comparing the log transformed fold change of cells in each cell type before and after perturbation.

### Markers, enrichments and differential expressions

Cell-type specific markers were identified by intersecting genes preferentially expressed in each cell type with TFs preferentially active in each cell type. Cell-type specific expression was obtained from comparing pseudobulk of each cell type against the other cell types, while cell-type specific TFs were derived by ranking TFs based on their outdegree in each cell-type specific GRN. Differential expression was performed using edgeR edg2 function^56^. Genes were selected using fold-change >= 2 and FDR <= 0.05 thresholds, calculating a score as -log10(FDR)*log2FC, and taking the top 20 expressed TFs in each cell type.

Gene Ontology enrichments were calculated using topGO 2.54 on R 4.3. HPO enrichments of a given gene set of interest were calculated as -log10(% of genes in the set of interest in a given HPO/size of the gene set of interest), and significance was calculated by hypergeometric test. The same approach was used to calculate enrichments for Excess (genes that accumulated an excess of high frequency mutations relative to their length in *Homo sapiens* with respect to other hominins), Length (genes showing increased length in *Homo sapiens*) and Select (genes associated with signatures of positive selection) lists from Kuhlwilm and Boeckx^21^.

---

**Supplementary Figure 1.**
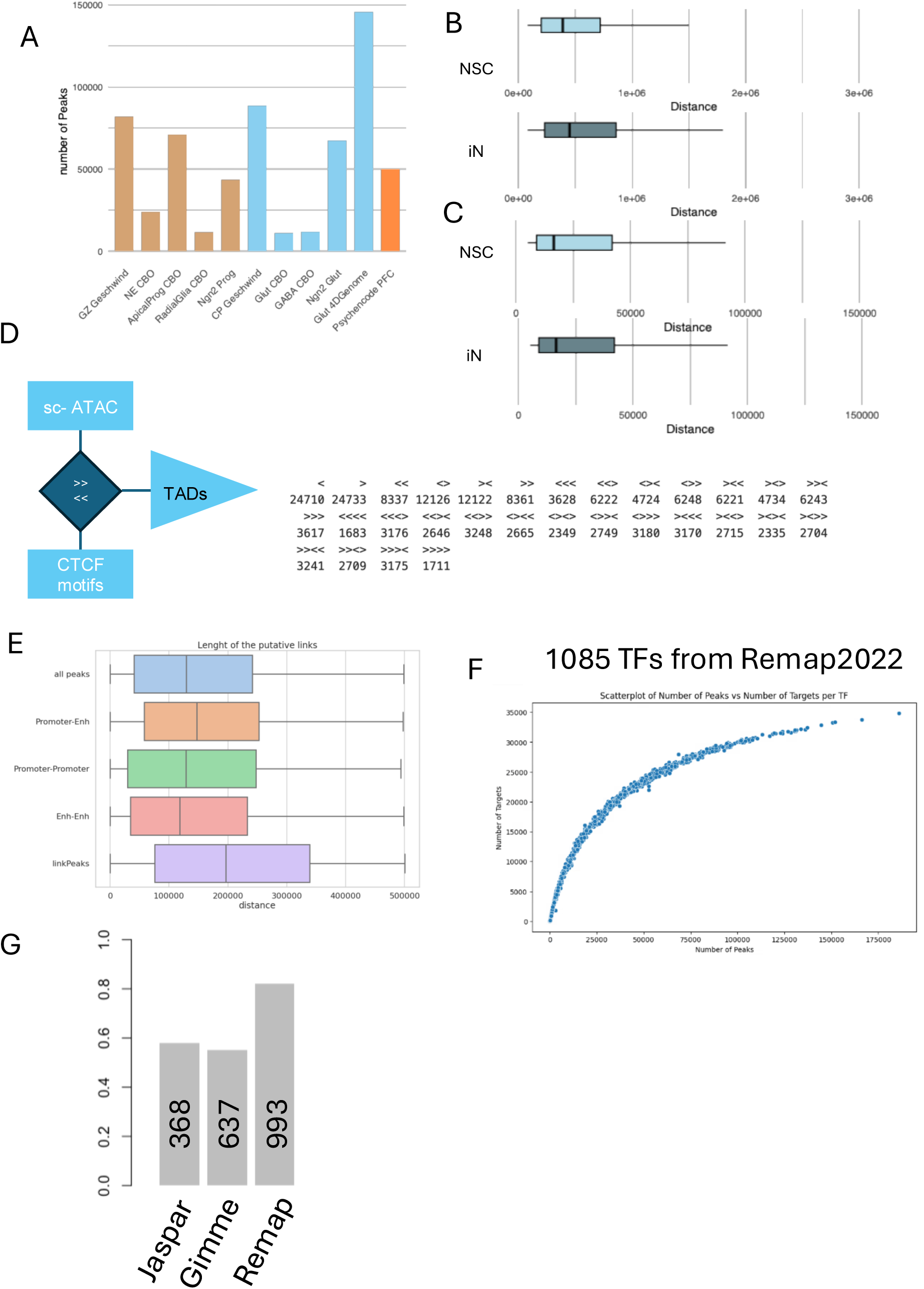
Characterization and validation of the CatReg reference. **A.** Bar plot showing the number of regulatory peaks contributed by each source dataset used to build CatReg. Germinal zone (GZ) and cortical plate (CP) H3K27ac peaks were derived from ref. ^12^; cortical brain organoids (CBO), Ngn2 progenitors (“Ngn2 Prog”), and Ngn2-induced neurons (“Ngn2 Glut”) from ref. ^13,40^; cortical glutamatergic neuron peaks (“Glut 4DGenome”) from ref. ^41^. “PsychENCODE PFC” corresponds to PsychENCODE-defined H3K27ac enhancer sites (DER-04a_hg38lft_PEC_enhancers). **B.** Number of RADICL-seq contacts in iPSCs, neural stem cells (NSCs) and neurons from the FANTOM6 dataset. **C.** Number of Hi-C contacts in iPSCs, NSCs and neurons from the FANTOM6 dataset. **D.** Number of CUT&Tag peaks for histone marks (H3K27ac, H3K27me3, H3K4me3 and H3K4me1) in neurons (FANTOM6), shown for two replicates. **E.** Boxplots of genomic distances spanned by Hi-C contacts across the FANTOM6 neurogenic series (NSC and induced neurons, iN), illustrating the broad distance distribution of Hi-C interactions. **F.** Boxplots of genomic distances spanned by RADICL-seq interactions across the FANTOM6 neurogenic series (NSC and iN), showing a shorter-range distribution relative to Hi-C (after filtering very short-range events, as described in Methods). **G.** Overview of the TADboundaries workflow (https://github.com/veronica-fin/TADboundaries) used to infer contact-like constraints from single-cell ATAC-seq co-accessibility combined with CTCF motif organization. The schematic summarizes how co-accessible peak pairs are filtered/prioritized using predicted structural features, with example CTCF motif orientation patterns indicated (comparative symbols represent motif directionality). **H.** Boxplots comparing distances of cis co-accessibility links included in CatReg (“all peaks”) and stratified by link type (promoter–enhancer, promoter–promoter, enhancer–enhancer), alongside distances reported by Seurat/Signac LinkPeaks. This panel highlights the shorter-range distribution of CatReg co-accessible links relative to LinkPeaks. **I.** Scatter plot relating TF occupancy to inferred regulatory output after intersecting ReMap ChIP-seq with CatReg. Each point is a TF; the x-axis indicates the number of CatReg peaks overlapped by that TF’s ReMap peak set, and the y-axis indicates the number of associated TF–gene targets recovered after intersection with CatReg, revealing a non-linear relationship consistent with enrichment for functional TF–gene links. **J.** Bar plot summarizing the fraction of CatReg targets supported by each TF/motif reference: JASPAR2024, GimmeMotifs (vertebrate), and ReMap.

**Supplementary Figure 2.**
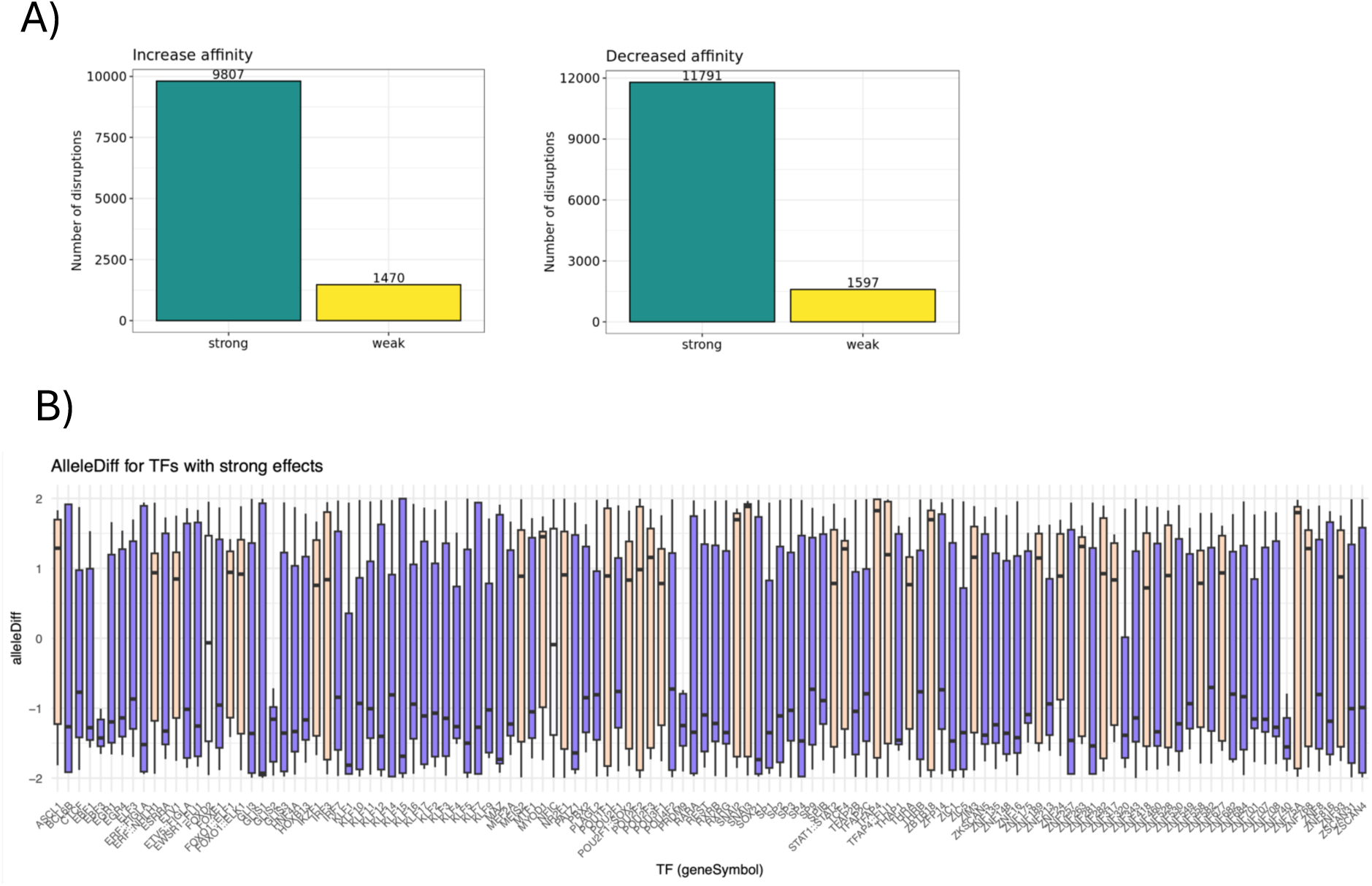
Predicted impact of human-specific high-frequency variants on TF binding motifs (MotifbreakR) (A) MotifbreakR summary of the predicted effect of human-specific high-frequency variants (HFVs) on transcription factor (TF) binding affinity. Bar plots show the number of HFV–motif disruptions predicted to increase (left) or decrease (right) motif binding affinity, stratified by effect size (strong vs weak) as reported by MotifbreakR. (B) Distribution of MotifbreakR alleleDiff scores for TFs with strong predicted effects. Boxplots summarize, for each TF motif, the allele-dependent change in predicted binding (alternate vs reference allele), highlighting TF-specific variability in the magnitude and direction of HFV-driven motif changes.

**Supplementary Figure 3.**
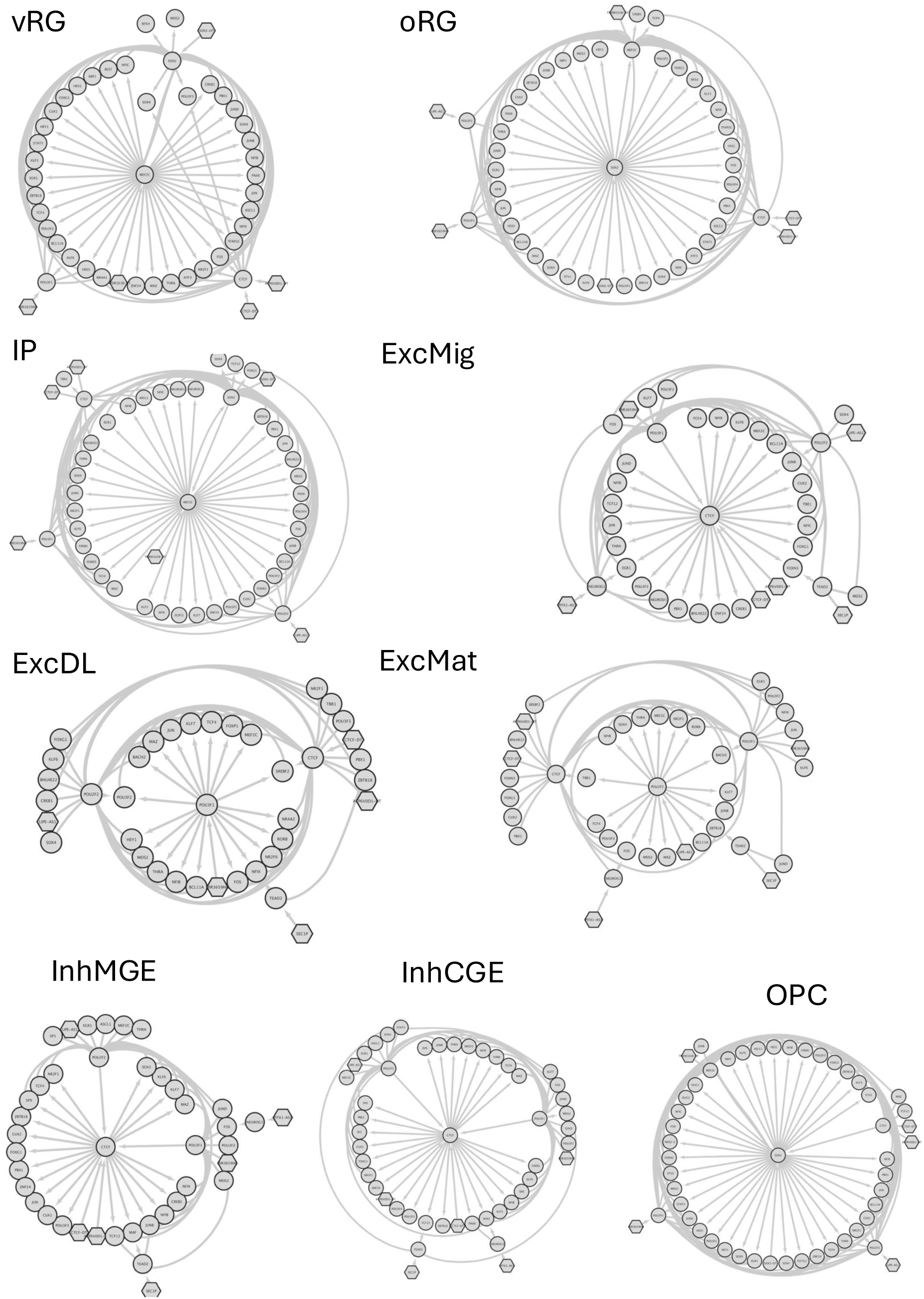
Cell type–resolved TF–ncRNA regulatory modules supported by GRNs and RADICL-seq. Network diagrams summarizing minimal TF–ncRNA modules for each neuronal lineage/state (vRG, oRG, IP, ExcMig, ExcDL, ExcMat, InhMGE, InhCGE, OPC). Nodes represent regulators whose connections are supported by gene regulatory networks (GRNs) and RNA–DNA proximity interactions (RADICL-seq). All transcription factors (TFs) shown are drawn from the set whose regulatory regions display modern human–specific differential motif affinity (MotifbreakR). Non-coding RNAs (ncRNAs) are included when they share at least two target genes with TFs in the same module. Circles denote TFs and hexagons denote ncRNAs; edges indicate shared targets and/or RADICL-supported associations within each cell type. Across lineages, TEAD2 forms cell type–specific partnerships with distinct ncRNAs, while LIPE-AS1 and SEC1P appear recurrently across multiple networks. The relative placement of neurogenin (NEUROG) genes varies between cell types, consistent with context-dependent positioning upstream or downstream of immediate-early regulators such as FOS.

**Supplementary Figure 4.**
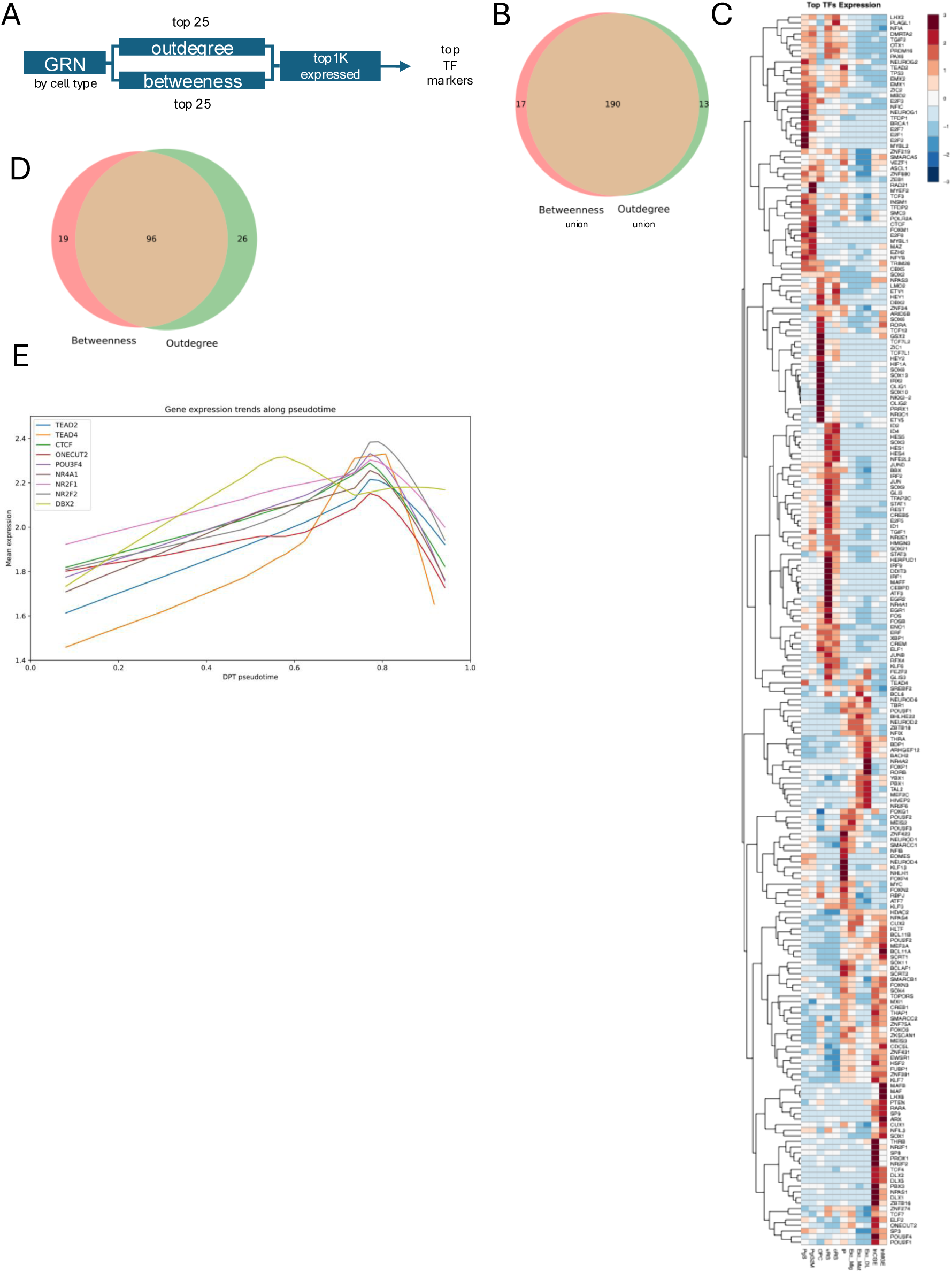
Prioritization of cell type–specific TF regulators and their expression dynamics. (A) Workflow used to prioritize transcription factors (TFs) by combining GRN connectivity and cell type–specific expression. For each cell type, TFs were ranked by network centrality (top hubs by outdegree and by betweenness) and intersected with highly expressed TFs (top 1,000 expressed genes per cell type) to define candidate TF markers/regulators. (B) Venn diagram showing the overlap between TFs prioritized as hubs when using the union across cell types of the top regulators ranked by betweenness versus the union of top regulators ranked by outdegree (default cutoff used in the main analysis). (C) Heatmap of normalized expression for the prioritized top TF markers across the neurogenic cell types/states, with hierarchical clustering highlighting shared and cell type–restricted TF expression programs. (D) Sensitivity analysis of hub selection: Venn diagram showing overlap between hub TFs identified by betweenness and outdegree when using a more stringent cutoff (top 25 per cell type, instead of the main cutoff). (E) Expression trajectories of selected TFs across differentiation pseudotime. Lines show smoothed expression trends for key TFs (HFV-associated and participating in TF–ncRNA modules), which peak early along the trajectory—consistent with higher activity in early/intermediate populations (e.g., IP / lower-layer excitatory / migrating excitatory states).

**Supplementary Figure 5.**
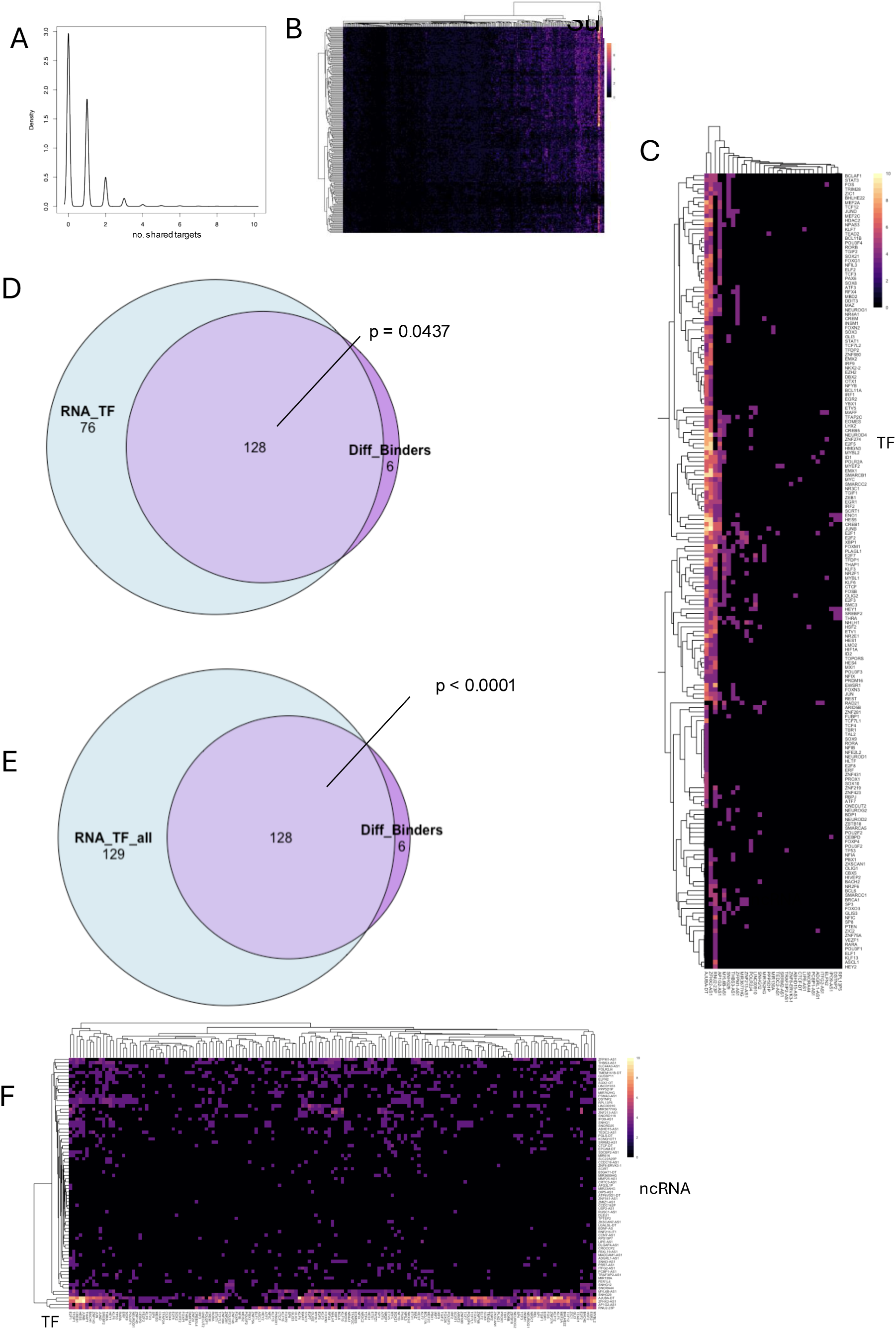
Shared-target links between RADICL-seq ncRNAs and GRN TFs and enrichment among HFV differential binders. (A) Density distribution of the number of shared target genes between ncRNAs detected by RADICL-seq and TF regulators inferred from cell type–resolved GRNs. Each ncRNA–TF pair is scored by the size of its shared target set. (B) Heatmap of shared-target counts for all ncRNA–TF pairs from (A), restricted to cell type–specific TF markers/regulators. Rows/columns are hierarchically clustered to highlight recurrent co-regulatory structure across TFs and ncRNAs. (C) Heatmap of the strongest ncRNA–TF associations, showing only pairs with ≥4 shared targets, emphasizing high-confidence co-regulatory modules. (D) Venn diagram testing overlap between RNA-co-regulating TF markers (TF markers that share targets with ≥1 RADICL ncRNA) and TFs predicted by MotifbreakR to show differential binding affinity driven by human-specific high-frequency variants (HFVs). The reported p value indicates enrichment of differential binders within the RNA-co-regulating TF marker set. (E) Same overlap analysis as in (D), but using the broader set of all RNA-co-regulating TFs (not limited to TF markers). The reported p value indicates significant enrichment of MotifbreakR differential binders among TFs that co-regulate targets with RADICL ncRNAs. (F) Heatmap restricted to TFs with MotifbreakR-predicted differential binding affinity that also participate in ncRNA shared-target relationships. While most ncRNA–TF pairs share few or no targets, a subset shows high shared-target counts (up to ∼10), highlighting candidate HFV-sensitive TF–ncRNA co-regulatory interactions.

